# Aberrant CD4^+^ T cell refeeding response impairs neuro-immune crosstalk in Parkinson’s disease

**DOI:** 10.64898/2026.05.06.723248

**Authors:** Karl Austin-Muttitt, Bethan R. David, Martina Sassi, Ravi Agarwal, Thais S. Sabedot, Jeffrey R. Jones, Michael S. Cuoco, Benjamin J. Jenkins, Luke D. Roberts, Leo Harris, Natalia Kieronczyk, Gang Han, Matthew D. Hitchings, Paul C. Donaghy, Alison J. Yarnall, Catherine A. Thornton, Biju Mohamed, Alwena H. Morgan, Ffion Thomas, Nicholas Jones, Fred H. Gage, Jeffrey S. Davies

## Abstract

T cells modulate disease associated neuroinflammation in Parkinson’s disease (PD). We report that circulating CD4^+^ T cells from people with PD have a dysregulated transcriptional and cytokine response to fasting and refeeding and an altered metabolic profile. The CD4^+^ T cell secretome mediates a metabolic program in neurons that is impaired in PD, revealing dysfunctional neuro-immune signalling that may contribute to disease pathology.

## Main

Parkinson’s disease (PD) is a neurodegenerative disorder characterized by motor symptoms but is also associated with significant metabolic and immune system dysregulation (Tansey, et al., 2022). Recent studies implicate autoimmune T cell responses to self-antigens in PD (Williams, et al., 2024). Notably, CD4^+^ T cells migrate from the blood to the midbrain and contribute to α-synuclein-mediated loss of dopaminergic neurons in rodent models of PD (Williams, et al., 2021; Harms, et al., 2017). Similarly, Th17 cells induce neuronal death in human induced pluripotent stem cell models of PD (Sommer, et al., 2018). Metabolic shifts in T cells, including increased glycolysis and mitochondrial dysfunction, are associated with increased pro-inflammatory signalling in the prodromal phase of PD (Mark, et al., 2025; Smith, et al., 2018). Immune cells undergo metabolic adaptation in response to low energy, notably via the mTOR/AMPK nutrient-sensing pathway, to maintain functionality (Blagih, et al., 2015; Pearce, et al., 2013), and whole body energy status modulates immune system function, with fasting linked to attenuated inflammation (Meydani, et al., 2016) and decreased immune cell counts in blood and lymphoid tissues (Contreras, et al., 2018). However, the interplay between metabolism and immune function in the context of PD remains relatively underexplored.

To validate the presence of T cells in the human brain, we quantified CD3^+^ cells in post mortem brain tissue from PD and control donors (Fig.1a). Analysis of substantia nigra (SN) tissue revealed a 1.7 fold increase in CD3^+^ cells in PD brains compared to control brains, suggesting increased T cell recruitment to the SN in PD (Fig. 1b,c). This is consistent with reports of increased CD4^+^ T cell infiltration in PD correlating with α-synuclein and tau aggregation (Kouli, et al., 2020). Furthermore, we quantified expression of two characteristic Th17 cell genes, IL17A and RORC, using *in situ* hybridization: expression of these transcripts was elevated by 2.3 and 2.1 fold respectively in the SN of PD donors (Fig.1d,e), supporting a role for Th17 cells in PD.

**Fig. 1:**
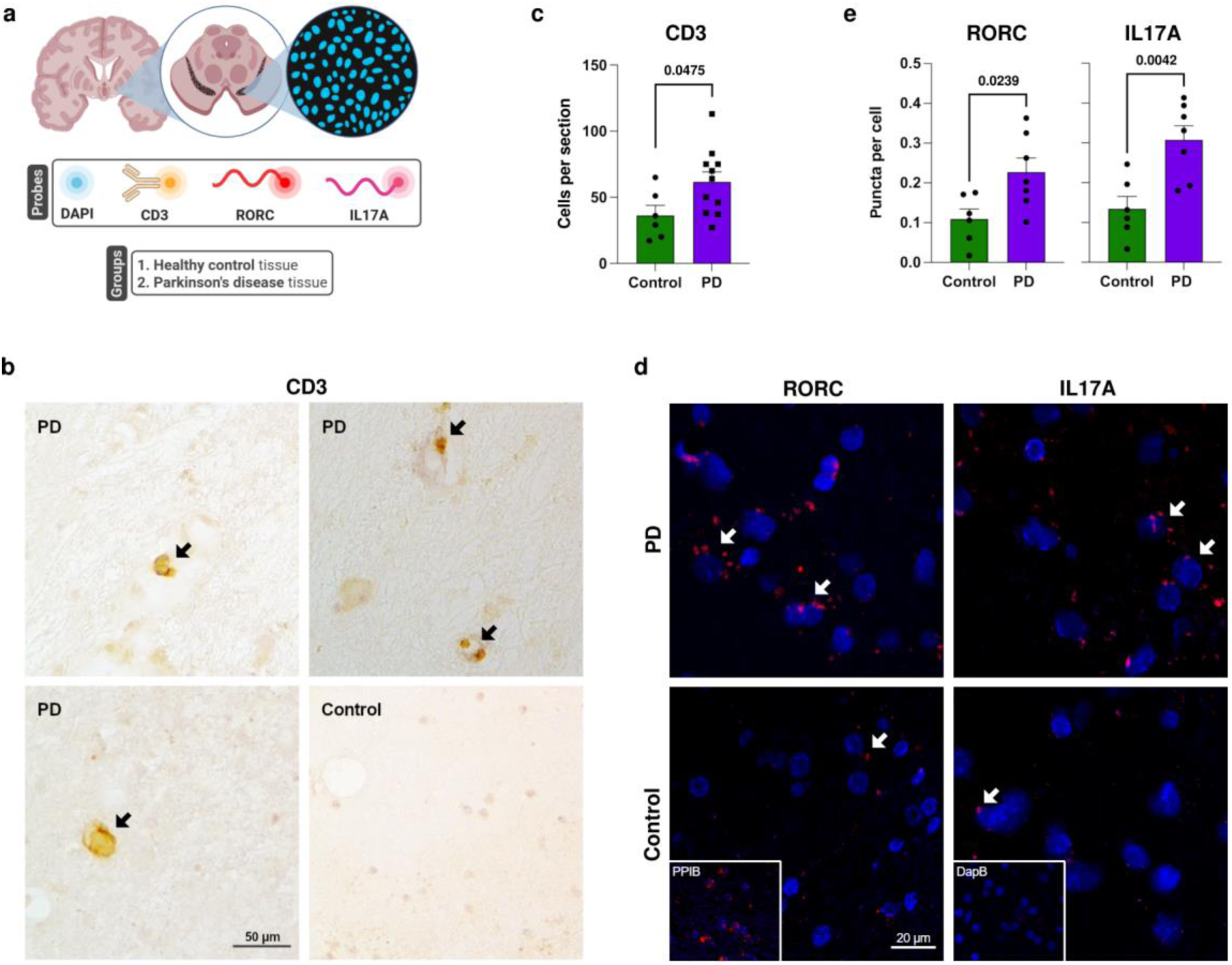
Evidence for increased infiltration of Th17 cells in the PD SN. **a**, Experiment schematic. **b**, Representative images of CD3^+^ cells in SN tissue sections. **c**, Comparing SN tissue from healthy control donors (*n* = 6) and PD donors (*n* = 11), there is a significant increase in CD3^+^ cells per section in the PD tissue (1.7 fold change; *p* = 0.0475); Student’s *t* test, *p* < 0.05. **d**, Representative images of IL17A and RORC transcripts in SN tissue from healthy control donors (*n* = 6) and PD donors (*n* = 7), measured via *in situ* hybridization, alongside DAPI staining of cell nuclei; PPIB positive control, bacterial DapB negative control. **e**, There are significant increases to RORC (2.1 fold change; *p* = 0.0239) and IL17A (2.3 fold change; p = 0.0042) puncta per cell; Student’s *t* test, *p* < 0.05.

Recent studies suggest a link between CD4^+^ T cell metabolism and impaired neuron function in PD (Tansey, et al., 2022), but as of yet there is no direct evidence of this. To address this gap in knowledge, we characterized the response of circulating CD4^+^ T cells from PD donors and age- and sex-matched controls to a fasting-refeeding paradigm (ClinicalTrials.gov ID: NCT05381090). Blood samples were collected following an overnight fast and 3 hours after refeeding (Fig.2a). Selected timepoints were chosen for their established responses to fasting in healthy individuals (Han, et al., 2021). Flow cytometry analysis identified a refeeding-mediated increase in total circulating CD4^+^ T cells in both control and PD donors (1.7 and 1.4 fold change, respectively), consistent with homeostatic fasting-induced T cell homing to bone marrow (Collins, et al., 2019) (Supplementary Fig.1). Next, we isolated CD4^+^ T cells from both the fasted and refed timepoints and stimulated them with anti-CD3/CD28 for 72 hours prior to multiplex analysis of cytokine secretion. From control donors, we observed the expected increase (1.7 fold change) in overall cytokine secretion after refeeding following an overnight fast. Strikingly, this response was not observed in CD4^+^ T cells from PD donors, suggesting a disease-associated impairment in cytokine secretion following metabolic challenge (Fig.2b,c). Remarkably, both the pro-inflammatory cytokine IFNγ and the anti-inflammatory cytokine IL-10 increased significantly in healthy control but not PD CD4^+^ T cells (Fig.2d).

In parallel, we performed RNA sequencing of isolated CD4^+^ T cells to assess their transcriptomic response. Our experimental design of paired fasting and refeeding samples revealed substantial differentially expressed genes (DEGs) associated with metabolic challenge in controls (413 genes). Gene ontology (GO) analysis of control DEGs (Fig.2c) showed substantial changes across the nucleosome, mitochondria, proteasome, and T cell fate. The transcription factor BRD4, reported to drive altered CD4^+^ T cell differentiation towards proinflammatory (Cheung, Zhang, & Jaganathan, et al., 2017) and neuroinflammatory (Dey, et al., 2025) immunophenotypes, was upregulated following refeeding. In contrast, no genes were significantly differentially expressed in CD4^+^ T cells from PD donors, again suggesting an impaired response to metabolic challenge (Fig.2c).

Modelling of transcription factor activity in control donor CD4^+^ T cells (Fig.2h) predicted, for example, that the mediator of the integrated stress response, ATF4, essential for immunosuppression in response to nutrient starvation in CD4^+^ T cells (Zou, et al., 2024), has downregulated activity. AHR, the transcription factor most strongly upregulated in the refed condition, promotes the differentiation of self-regulatory IFNγ^+^ IL-10^+^ Tr1 cells (Apetoh, et al., 2010). GO analysis identified increased canonical histone content consistent with enhanced proliferation supported by the upregulation of the G1/S-transition-associated transcription factor, TFDP1. TGFB1 signalling and RUNX1 activity, also upregulated, are fate-determining pathways in CD4^+^ T cells that are attenuated by AMPK signalling (Yin, et al., 2023; Gayatri, et al., 2023). While control donor CD4^+^ T cells showed altered activation of diverse pathways following refeeding, these changes were absent in PD donor CD4^+^ T cells.

Mitochondria emerged as a key node in the fasting-refeeding response in control donor CD4^+^ T cells, with GO analysis revealing downregulation of both oxidative phosphorylation and mitochondrial autophagy in the refed condition (Fig.2f,g). The PD-associated redox sensor PARK7 (encoding DJ-1) and redox control enzyme NQO1 are downregulated alongside oxidative phosphorylation, as are key mitochondrial quality control genes (e.g. GABARAP, GABARAPL2, NIPSNAP3A, PARL). These pathways were not significantly regulated in PD.

To further assess metabolic responses in CD4^+^ T cells, we analyzed the transcriptome data using Compass, a flux balance analysis tool developed to predict metabolic state in Th17 CD4^+^ T cells (Wagner, et al., 2021). In control donors, the refed cells exhibited decreased mitochondrial transport (mean *d* = −2.3) and fatty acid oxidation (mean *d* = −1.7; Fig.2i). We also observed the upregulation of key genes for fatty acid synthesis (SREBF1, FASN, FADS1, SCD), reflecting a program of metabolic rewiring associated with pro-inflammatory activity in T cells (Hu, et al., 2024). In stark contrast to control donors, the analysis predicted increased mitochondrial transport and fatty acid oxidation in refed PD cells (mean *d* = 1.4 and 1.3, respectively), suggesting profound PD-linked dysregulation in the fasting-refeeding response.

These findings identify DEGs and associated pathways in control donor CD4^+^ T cells that putatively represent the homeostatic response to metabolic challenge. Importantly, analysis of these same fasting-refeeding responses in CD4^+^ T cells from PD donors revealed profound metabolic inanition consistent with mitochondrial deficits reported in SN dopaminergic neurons that are susceptible to damage in PD (Schapira, et al., 1989).

Next, as CD4^+^ T cells migrate to the brain and exacerbate neuropathology (Brochard, et al., 2009; Tansey, et al., 2022), we asked whether the fasting-refeeding responses as described above signal metabolic and inflammatory state to neurons. To do this, we established a cell-cell communication assay whereby induced neurons (iNs) were exposed to the secretome of CD4^+^ T cells. Briefly, fibroblasts collected from healthy control or sporadic PD donors underwent neuron conversion for 21 days. From day 20, neurons were cultured for the remaining 24 hours in pooled supernatant collected from disease-matched CD4^+^ T cells under either a fasted or refed state, treating control iNs with control supernatants and PD iNs with PD supernatants (Fig.3a; Supplementary Table 3). Importantly, iNs retain age-related epigenetic signatures underpinning neuronal impairments in neurodegenerative pathology (Mertens, et al., 2015; Mertens, et al., 2021).

A multi-omic mass spectrometry analysis of the pooled supernatants suggested robust differences in content between the fasted and refed pooled supernatant treatments in healthy controls, differences that were distinct and blunted, but not abolished, in PD (Supplementary Fig.2). In both healthy control and PD iN lines, we found a stark transcriptomic response to vehicle media treatment (Supplementary Fig.3), confirming that healthy control and PD iNs are responsive to external stimuli. We next compared the iN transcriptomic response to supernatant treatments (Fig.3a). In healthy control iNs, 1597 DEGs (Fig.3b) were subject to an overrepresentation analysis of GO terms that showed wide-ranging changes between supernatant treatments (Fig.3c,d). Broadly, pathways governing neuronal morphology and synaptic function were decreased in the refed treatment, whereas pathways for stress, inflammation, and neuro-immune crosstalk were enhanced. Investigation of these terms revealed downregulated axonal capacity (e.g. NEFL, MAPT), alongside genes relating to action potential (e.g. KCNA2, SCN8A, CACNA1H) and presynaptic neurotransmitter release (e.g. SYT7, SYT13, SV2A, RAB3A, STXBP1). This differential response to fasted and refed CD4^+^ T cell secretome treatment was absent in PD iNs.

**Fig. 2:**
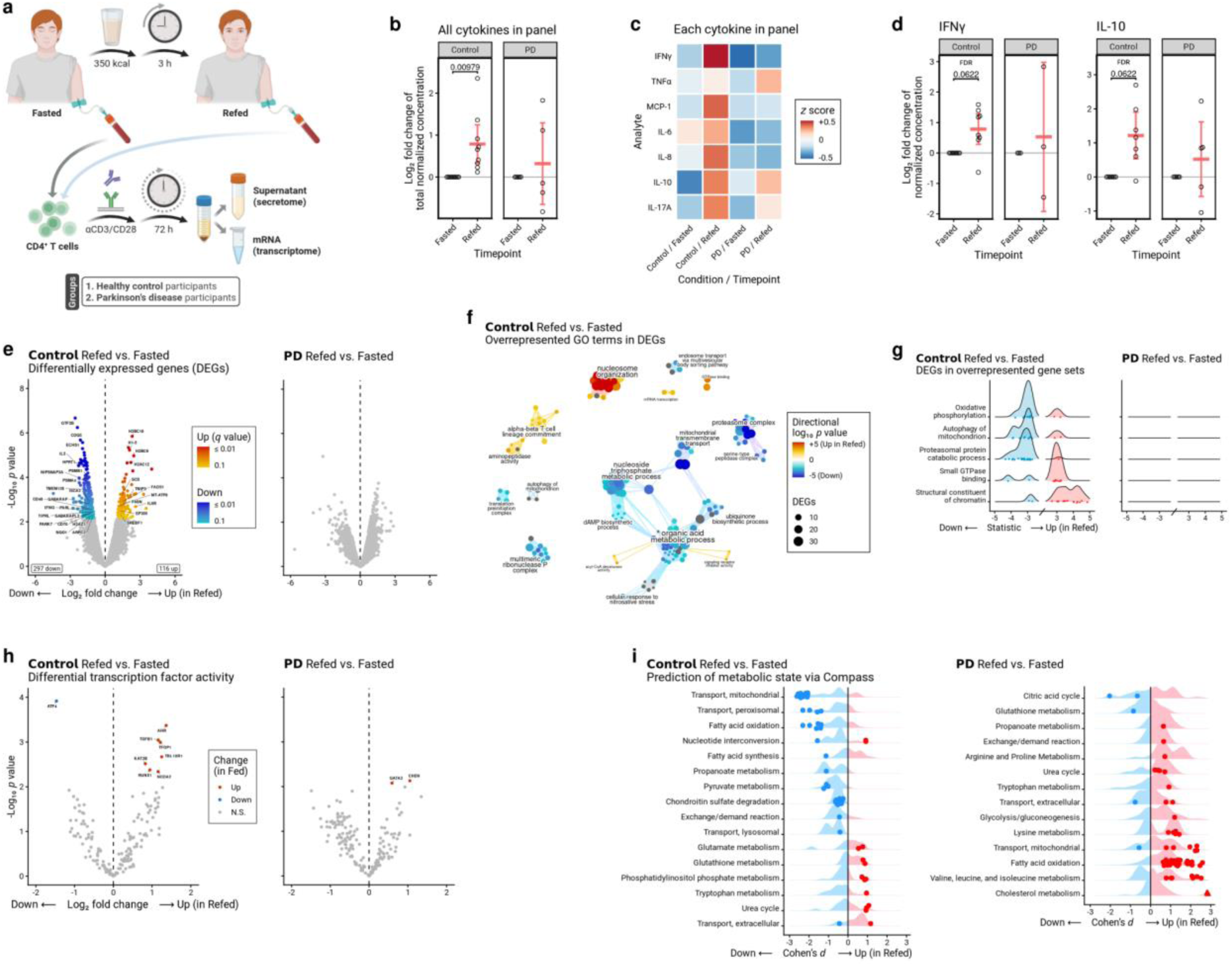
Fasting-refeeding response in CD4^+^ T cells is dysregulated in PD. **a**, Experiment schematic. **b**, Total normalised concentration of cytokines across panel measured in refed relative to fasted in healthy control donor CD4^+^ T cells (*n* = 9) and PD (*n* = 5), showing a significant increase in secretion from healthy control cells (*p* = 0.00979); paired t tests, *p* < 0.05. **c**, Heatmap illustrating the pattern of cytokine secretion across all experimental groups; mean normalised concentration transformed to z scores within each cytokine. **d**, Secretion of IFNγ and IL-10 measured in refed relative to fasted, showing a significant increase in secretion from healthy control cells (IFNγ: *FDR* = 0.0622; IL-10: *FDR* = 0.0622) but not PD cells; paired *t* tests adjusted with the Benjamini-Hochberg FDR procedure, *FDR* < 0.1. **e**, Differential gene expression analysis showing 413 DEGs when comparing refed to fasted healthy control donor cells (*n* = 5), and no DEGs in PD (*n* = 5); edgeR, likelihood ratio tests adjusted with Storey’s procedure, *q* < 0.1. **f**, Pathway cluster analysis of overrepresented GO terms identifies key themes in the response of healthy control cells: changes to nucleosomes, mitochondria, proteasomes, metabolism, and altered T cell immunophenotype; Limma (*goana*) and aPEAR, one sided hypergeometric tests, *p* < 0.01. **g**, Gene expression changes in highlighted GO terms; the statistic is the signed root likelihood ratio. **h**, Differences in modelled transcription factor activity scores indicating that the transcriptional program driving the fasting-refeeding response is altered between healthy control cells and PD cells; Priori, paired *t* tests, *p* < 0.01. **i**, Flux balance analysis of metabolic activity comparing reaction consistency scores between the refed and fasted cells, showing categories with at least one significant reaction, points are significant reactions, triangles capped for plotting; Compass, paired *t* tests, *p* < 0.01.

**Fig. 3:**
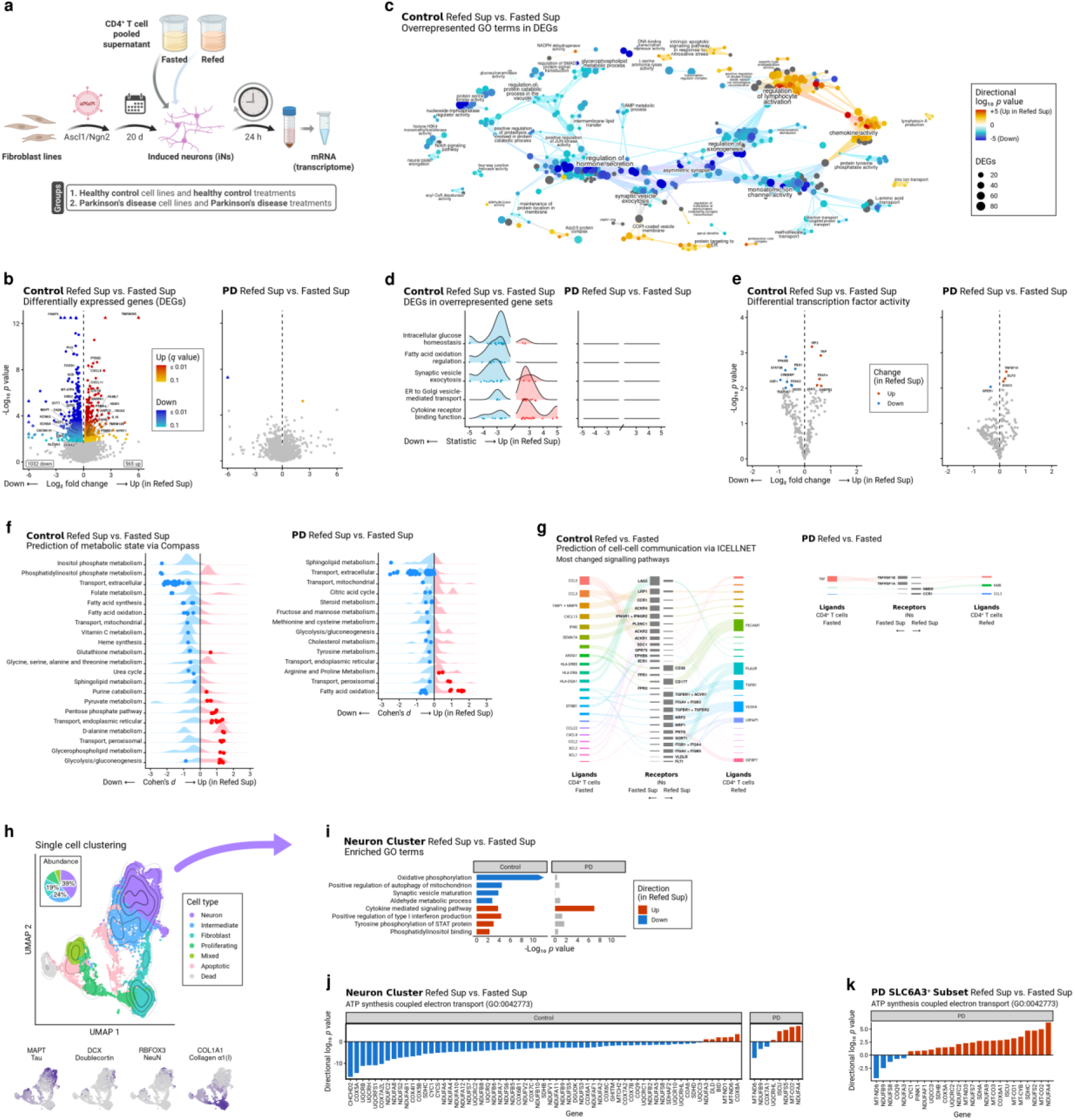
Regulation of neuron metabolism in response to fasted-refed CD4^+^ T cell secretomes is dysfunctional in PD. **a**, Experiment schematic. **b**, Differential gene expression analysis showing 1597 DEGs when comparing refed CD4^+^ T cell supernatant treatment to fasted treatment in healthy control iNs (*n* = 4) and 2 DEGs in PD iNs (*n* = 4), triangles capped for plotting; edgeR, likelihood ratio tests adjusted with Storey’s procedure, *q* < 0.1. **c**, Pathway cluster analysis of overrepresented GO terms identifies key themes in the differential response of healthy control iNs to treatment: changes to axonal morphology, synaptic vesicle secretion, metabolism, and cytokine and chemokine release; Limma (*goana*) and aPEAR, one sided hypergeometric tests, *p* < 0.01. **d**, Gene expression changes in highlighted GO terms; the statistic is the signed root likelihood ratio. **e**, Differences in modelled transcription factor activity scores indicating that the transcriptional program driving the response to treatment with fasted-refed CD4^+^ T cell secretomes is altered between healthy control iNs and PD iNs; Priori, paired *t* tests, *p* < 0.01. **f**, Flux balance analysis of metabolic activity comparing reaction consistency scores between the refed and fasted treatments, showing categories with at least one significant reaction, points are significant reactions; Compass, paired *t* tests, p < 0.01. **g**, Modelled cell-cell communication from the CD4^+^ T cells to the iNs, where the signalling pathways shown are those with a large change in activity score between treatments, |log_2_ FC| > 3; ICELLNET. **h**, Dimensionality reduction and clustering of single cells (healthy control: *n* = 3605; PD: *n* = 5754), Seurat; left, marker genes; right, clusters; inset right: cell type proportions of all cells. **i**, Enrichment analysis of GO terms; Limma (*cameraPR*), Student’s *t* test allowing for correlation, *p* < 0.01. **j**, Gene expression in ATP synthesis coupled electron transport (GO:0042773). **k**, PD SLC6A3^+^ neuron gene expression in ATP synthesis coupled electron transport (GO:0042773).

When investigating genes essential to the dopaminergic neuronal phenotype in the healthy control iNs, we found downregulation of the core dopaminergic identity (SLC6A3, DRD2) as well as of FOXA2, a master regulator transcription factor associated with dopaminergic neuron survival under stress (Kittappa, et al., 2007) (Fig.3b). We observed upregulation of PD-associated redox control genes (PARK7, NQO1), also regulated in our control donor CD4^+^ T cells above. FKBP5, a PINK1-modulated driver of neuronal stress responses (Boonying, et al., 2019) reported in peripheral cells to be a fasting induced regulator (Han, et al., 2021), was downregulated. Additionally, the control iNs expressed immune signalling factors in response to the refed supernatant treatment: canonical IFNG-induced chemokines were upregulated (CXCL9, CXCL11) alongside IL10, a program suggestive of anti-inflammatory feedback to the immune compartment. In contrast, these pathways were not differentially expressed in iNs from PD donors, demonstrating impaired neuro-immune crosstalk under these conditions (Fig.3b,d).

Transcription factor activity analysis in control iNs (Fig.3e) offered further insight into the drivers of these phenotypic changes. The refed supernatant treatment remodels JAK-STAT expression away from STAT5B and towards JAK2, which shifts gene expression from neurotrophic factors towards neuroinflammation (Byts, et al., 2008; Wang, et al., 2019). The increased activity of IRF2 is consistent with a negative feedback mechanism to control IFNG driven expression (Lukhele, et al., 2022).

We found evidence that factors secreted from healthy control CD4^+^ T cells following metabolic challenge signalled healthy control iNs to engage an altered program of metabolism. Genes with the glucose homeostasis ontology were strongly downregulated. Flux balance analysis (Fig.3f) supported a Warburg-like switch in the metabolic character of the iNs, predicting enhanced glycolysis following refed treatment (mean *d* = 0.9), with reduced fatty acid oxidation (mean *d* = −0.8) and fatty acid synthesis (mean *d* = −0.7; DEGs: FASN ↓, FADS1 ↓, SCD ↓). In contrast, the PD iNs did not show Warburg-like switching following treatment with refed supernatant, identifying aberrant neuro-immune metabolic crosstalk in PD.

A cell-cell communication analysis (Fig.3g) cross-referenced ligand expression in the CD4^+^ T cell transcriptome with receptor expression in the iN transcriptome. In healthy controls, it predicted wide-ranging changes to signalling patterns between T cells and iNs in the refed condition versus the fasted condition, including to the canonical IFNG and TGFB1 pathways, as well as to the CCR1 pro-inflammatory chemokine signalling pathway (Yan, et al., 2020). In PD, only one substantive differential signal was predicted: a modest change to TNF signalling.

As transdifferentiation results in a mixed population of cells, single cell sequencing analysis was performed to confirm the effect of the fasted and refed treatments in the subpopulation of iNs that made up the MAPT^high^ COL1A1^low^ neuronal cluster (Fig.3h). In the healthy control neuronal cluster, we confirmed that treatment with refed supernatant induced a Warburg-like metabolic switch, decreasing oxidative phosphorylation as part of a program of metabolic and signalling changes (Fig.3i). In contrast, while the PD neuronal cluster responded to cytokine stimulus, it failed to regulate oxidative phosphorylation or to mount the wider changes to metabolism. This prompted us to analyze changes to the expression of individual genes involved with ATP-synthesis-coupled electron transport (Fig.3j). In healthy controls we found that transcription was strongly downregulated with the refed treatment, with 57 genes decreased. Notably, two of the most strongly downregulated genes were the PD-associated Complex I subunit NDUFS2 (Gonzalez-Rodriguez, et al., 2021) and mitochondrial regulatory factor CHCHD2 (Funayama, et al., 2015). Conversely, in the PD neuronal cluster, these genes were not regulated.

Dopaminergic neurons exhibit particularly high metabolic demand, which is implicated in their vulnerability in PD (Pacelli, et al., 2015). In light of our observation about the loss of dopaminergic capacity between fasting and refeeding (Fig.3b), we hypothesized that PD dopamine-transporter-positive (SLC6A3^+^) iNs would have more strongly dysfunctional expression of ATP-synthesis-coupled electron transport machinery. We analyzed the marker genes between treatments for this SLC6A3^+^ subset and confirmed that these neurons had dysfunctional upregulation, with 21 upregulated genes (Fig.3k), which was not observed in the healthy control iN neuronal cluster.

In summary, our data suggest convergence of PD pathways linked to energy balance in both human CD4^+^ T cells and neurons. We provide evidence for neuro-immune crosstalk signalling metabolic challenge and show that this crosstalk is aberrant in PD. Notably, we uncover a new role for the peripheral immune system in PD that could lead to delineation of pathogenic mechanisms with immunotherapeutic relevance.

## Methods

### Human post-mortem brain tissue

Human brain tissue was obtained from the Parkinson’s UK Brain Bank at Imperial College London with ethical approval (07/MRE09/72). Post-mortem brain tissue sections included healthy controls (*n* = 6) with no indications of neurodegenerative disease and participants diagnosed with PD (*n* = 11). Formalin-fixed frozen tissue was cryo-sectioned to 6–8 μm thickness, placed onto SuperFrost+ coated slides (VWR), and stored at −70°C. All cases had a post-mortem interval of < 42 hours. (See Supplementary Table 1.)

### DAB immunohistochemistry

Healthy control (*n* = 6) and PD (*n* = 11) human brain substantia nigra (SN) tissue sections were removed from the freezer and air-dried for 1-3 minutes before a hydrophobic pen was used to create a barrier around each section. A series of three 10 minute PBS washes was performed, followed by incubation with 1% H_2_O_2_ (stock 30%) in 0.15% PBST for 8 minutes. Sections were washed again and antigen retrieval performed using citraconic anhydride buffer 98% (1:2000 in distilled water) for 30 minutes at 100 °C in the steamer. Sections were blocked with 10% normal goat serum (NGS) in 0.15% PBST for 20 minutes at room temperature, followed by incubation with polyclonal rabbit anti-human CD3 (Dako, Cat: A0452) diluted 1:500 in 0.15% PBST, overnight at 4 °C.

On the second day, sections were washed again and incubated with biotinylated goat anti-rabbit antibody (Vector Labs, Cat: BA-1000-1.5) diluted 1:500 in 0.15% PBST. Another series of three 10 minute washes was performed before incubation with ABC solution (VECTASTAIN ABC-HRP Kit, Vector Labs, Cat: PK-4000) for 60 minutes at RT. Another set of washes was followed by detection of immunoreactivity using ImmPACT DAB (Vector Labs, Cat: SK-4103) for 90 seconds. Sections were then placed in warm tap water, counterstained with haematoxylin for 90 seconds, and further destained in warm tap water. Dehydration (70% for 3 minutes and 100% ethanol for 5 minutes) and delipidation (4x Histoclear, 1 minute each) steps were performed before sections were coverslipped using Entellan mounting medium Sigma Aldrich).

CD3 immunoreactive sections were analysed using a Nikon Eclipse 50i microscope and CD3^+^ cells were manually counted using a 40x objective lens across the SN and expressed as cells per section. All analyses were performed blind to phenotype. Statistical analysis was performed using Student’s *t* test in GraphPad Prism 9.0.0, with *p* < 0.05 considered significant. Representative images were enhanced in contrast and labelled.

### In situ hybridization (ISH)

Two ISH analyses (RNAScope) were performed on the SN tissue present in post mortem brain sections from healthy control donors (*n* = 6) and PD donors (*n* = 7): the first with a probe targeting the transcript for IL-17A (HiPlex Probe Hs-IL-17A, Bio-Techne, Cat: 310931), and the second with a probe targeting the transcript for RORγt (HiPlex Probe Hs-RORGT, Bio-Techne, Cat: 311061). A positive control probe targeting the housekeeping gene PPIB (HiPlex Probe Hs-PPIB, Bio-Techne, Cat: 313901) was applied to tissues from at least one case per batch to confirm RNA integrity and to confirm consistency of expression. A probe targeting the bacterial DapB gene was used as negative control for each ISH run to exclude non-specific staining of the probes. All reagents — distilled water, ethanol, and wash buffers — were prepared fresh for each experiment.

The slides were allowed to thaw for 2–3 minutes at room temperature and then sequentially dehydrated in 70% to 100% ethanol (3 minutes each). Slides were then baked at 60°C for 30 minutes to ensure tissue adherence. Subsequently, 3–8 drops of hydrogen peroxide were applied directly to each tissue section for a 10 minute incubation period at room temperature.

Antigen retrieval was performed by first rinsing the slides in distilled water (3–5 gentle dips) and immersion in 100 mL of 1x target retrieval reagent (prepared by diluting 10 mL of 10x solution in molecular grade water) in a steamer for 15 minutes after boiling. After recovery, the slides were returned to distilled water and washed twice in 100% ethanol (3 minutes per wash) before drying in a 60 °C incubator for 10–20 minutes.

Dried sections were treated with 3–8 drops of protease IV at 40°C for 30 minutes after forming a hydrophobic barrier around each tissue section. Immediately after washing in distilled water (ensuring that the slides did not dry), the appropriate target probe was applied (3–8 drops), and the slides were incubated at 40 °C for 2 hours. After incubation, sections were stored overnight in 5x SSC buffer at room temperature.

The following day, amplification reagents (AMP0–AMP6) were equilibrated to room temperature for 30 minutes before use. The sections were washed twice in 1X wash buffer and then sequentially processed with the AMP reagents: AMP1 (30 minutes at 40 °C), AMP2 (15 minutes at 40 °C), AMP3 (30 minutes at 40 °C), AMP4 (15 minutes at 40 °C), AMP5 (30 minutes at room temperature) and AMP6 (15 minutes at room temperature). The slides were washed three times (5 minutes per wash) with 1x wash buffer between each incubation. Fast Red chromogen was prepared by diluting Fast Red-B with Fast Red-A (1:60) and sections were incubated with the staining solution for 10 minutes at room temperature. This reaction was stopped by rinsing in tap water. Finally, the sections were counterstained with DAPI for 5 minutes, rinsed thoroughly in PBS, and dried in a 60 °C oven for 45 minutes. Dried sections were mounted using VectaMount (Vector Labs) and imaged the following day.

Each image was analyzed using Qupath (Bankhead, et al., 2017). The cell detection function was applied to the Hoechst channel (Life Technologies) and the subcellular detection function was applied to the probe channels. Each image was analyzed at 20x magnification using the same parameters. Average puncta per cell was calculated based on 4 images per case, dividing the number of subcellular puncta by the number of nuclei for the corresponding image. Statistical testing was undertaken using GraphPad Prism 9.0.0 and differences in puncta per cell were tested for significance (Student’s *t* test, *p* < 0.05). Representative images were enhanced in contrast and labelled.

### Blood draws from participants when fasted and refed

The study involving humans was approved by Research Ethics Committee (REC reference: 21/WA/0049; IRAS ID: 288862; ClinicalTrials.gov ID: NCT05381090) and was in accordance with the ethical principles that originate from the Declaration of Helsinki and that are consistent with the International Council for Harmonisation Guidelines on Good Clinical Practice (ICH E6 GCP) and the Mental Capacity Act 2005, with Research Governance maintained by Swansea University. Blood was drawn from healthy control participants (*n* = 9) and PD participants (*n* = 12). Subjects stopped taking cholinesterase-inhibitor medications prior to blood draw (e.g Galantamine 24h prior to clinic and Rivastigmine patch 24 hours prior to clinic or Rivastigmine tablets 12h prior to clinic). After a 12 hour fast, a small venous cannula was inserted into a vein to collect blood samples, which was kept patent with a 0.9% saline flush. A small amount of blood (< 1 mL) was drawn prior to the sample to ensure that it was not contaminated with saline. A fasted blood sample was collected in four 9 mL lithium heparin tubes (VACUETTE LH, Greiner Bio-One, Cat: 455084) and processed within 10 minutes of collection.

Participants were then given a standard liquid meal to drink (Ensure Plus vanilla nutrition supplement drink; 237 mL, 350 calories). 3 hours following the liquid meal, refed blood samples were collected using the same method as for the fasted blood draw.

### Isolation of peripheral blood mononuclear cells (PBMCs) and CD4^+^ T cells

Mononuclear cells were initially isolated by layering whole blood (1:1) onto Lymphoprep (StemCell Technologies, Cat: 07851) before centrifugation at 805 x *g* for 20 minutes at room temperature. Mononuclear cells were removed and washed with RPMI 1640 (Life Technologies, Cat: 61870036) twice by centrifugation at 515 x *g*. Human CD4^+^ T cell subsets were isolated using autoMACS cell separation (Miltenyi Biotec) via negative selection (Human CD4^+^ T Cell Isolation Kit, Miltenyi Biotec, Cat: 130-096-533). CD4^+^ T cells were then activated with plate-bound anti-CD3 (2 μg/mL; OKT3, BioLegend, Cat: 317326) and free anti-CD28 (20 μg/mL; CD28.2, BioLegend, Cat: 302934) in phenol red free RPMI (Gibco, Cat: 10363083) and GlutaMAX supplement (Gibco, Cat: 11554516) at 37 °C in 5% CO_2_ for 72 hours. After the initial 3 hours, the media was supplemented with 10% fetal calf serum (Cytiva, Cat: 11531831) to enhance T cell activation. At 48 hours half the media was replaced with fresh media before the T cell supernatants were collected at 72 hours, and cells were pelleted by centrifugation (330 x g for 10 minutes at 4 °C).

### Abundance of PBMCs and CD4^+^ T cells

A 10 μL volume was removed from the sample and diluted in a 1:10 ratio with RPMI. Nonviable cells were dyed with an equal volume of 0.4% trypan blue (Countess Cell Counting Chamber Slides, Invitrogen, Cat: C10228). A brightfield cell counter (Countess 3 FL Automated Cell Counter, Invitrogen, Cat: 16832556) was then used to quantify the abundance of viable cells. CD4^+^ T cell purity was measured using FITC anti-CD3 (mIgG2a, clone OKT3, BioLegend, Cat: 317306) and Alexa Fluor 647 anti-CD4 (mIgG2b, clone OKT4, BioLegend, Cat: 317422) antibodies. The purity of the isolated T cells was > 95% for all samples.

### Cytokine secretion by CD4^+^ T cells

Control samples (*n* = 9) and a randomly selected subset of PD samples (*n* = 5) underwent cytokine analysis. The supernatant from each sample was assayed for a selection of inflammatory cytokines using a 13-plex panel (BioLegend LEGENDplex Human Inflammation Panel 1, Cat: 740809). Fluorescence intensity data were processed using the manufacturer’s suggested software workflow (LEGENDplex Data Analysis Software Suite 8; BioLegend) and concentrations converted to molar units. Samples were normalized to a common median molar concentration. For statistical analysis all measurements within the experimental limits of detection were included and a capped value was included for those above the upper limit. Differences were tested for significance (paired *t* test, *p* < 0.05). A paired comparison was not possible if both measurements were above the upper limit. When multiple comparisons were performed, *p* values were adjusted and considered significant when *FDR* < 0.1 (Benjamini-Hochberg procedure).

### Bulk sequencing of CD4^+^ T cells

RNA sequencing and analysis was performed for a subset of the control participants (*n* = 5) and PD participants (*n* = 5). RNA isolation was performed on the pelleted cells for each sample (Cytiva RNAspin Mini, Cat: 25-0500-72; Qiagen RNeasy Plus Mini Kit, Cat: 74134) and RNA concentration and integrity was quantified (Qubit RNA HS Assay, Cat: Q32852). For each sample, 200 ng of total RNA was used as input for poly(A)-selected cDNA library preparation. Libraries were prepared using Oxford Nanopore Technologies cDNA-PCR barcoding kits compatible with either R9.4.1 or R10.4.1 flow cells. Sequencing was performed on a PromethION P2 Solo platform (Oxford Nanopore Technologies) using PromethION R9.4.1 (FLO-PRO002) and R10.4.1 (FLO-PRO114) flow cells at Swansea University Medical School. Samples were distributed across chemistries without bias. Basecalling was performed using ONT Dorado (HAC model). Transcripts were trimmed to remove sequencing adaptors and poly(A) tails (Pychopper; Fastplong) and filtered to a minimum length of 150 nt and minimum quality score of *Q* ≥ 9 (FASTQ-Filter); they then underwent pseudoalignment (Oarfish) (Zare Jousheghani, et al., 2025) to the human transcriptome (GENCODE 48). Sample labels were cross-referenced with sex-specific gene markers and HLA variant typing (T1K) (Song, et al., 2023). In the statistical model, the proportion of reads successfully mapped to the transcriptome was included as a covariate to account for differing degrees of RNA quality between samples.

### Pooling of CD4^+^ T cell secretomes into treatments

The supernatant from CD4^+^ T cells cultured, as described above, were collected at 72 hours and frozen for subsequent use. Briefly, 200 µL of supernatant from each participant (control, *n* = 9; PD, *n* = 10) was pooled in a group (control, PD) and treatment (fasted, refed) dependent manner, and frozen. Supernatant underwent two freeze-thaw cycles prior to being cultured with iNs for 24 hours (see below). To determine the content of pooled supernatants we performed mass spectrometry analysis for proteins, lipids, and metabolites (see below). A qualitative analysis of the fasting-refeeding response was undertaken by categorising analytes into upregulated, downregulated, or no change based on a fold change threshold of *FC* > 1.5 in either direction.

### Direct conversion of adult human fibroblasts into iNs

Primary human dermal fibroblasts (as previously described in Mertens, et al., 2015), maintained by the Stem Core at the Salk Institute, were used throughout. Fibroblasts originated from aged healthy control donors (*n* = 4) and donors with PD (*n* = 4). Protocols were previously approved by the Salk Institute Institutional Review Board and informed consent was obtained from all subjects.

Cells grown in 6-well plates were cultured in DMEM containing 15% tetracycline-free fetal bovine serum and 0.1% NEAA (Life Technologies), transduced with lentiviral particles for EtO and XTP-Ngn2:2A:Ascl1 (N2A), and expanded in the presence of puromycin (1 µg/mL; Sigma Aldrich). For iN conversion, fibroblasts were cultured in high densities and after 24 hours the medium was changed to neuron conversion (NC) medium based on DMEM:F12/Neurobasal (1:1) for 21 days. NC contains the following supplements: N2 supplement, B27 supplement (both 1x; GIBCO), doxycycline (2 µg/mL; Sigma Aldrich), Laminin (1 µg/mL; Life Technologies), dibutyryl cAMP (500 µg/mL; Sigma Aldrich), human recombinant noggin (150 ng/mL; Preprotech), LDN-193189 (0.5 µM; Cayman Chemicals) and A83-1 (0.5 µM; Stemgent), CHIR99021 (3 µM; LC Laboratories), and SB-431542 (10 µM; Cayman Chemicals). Medium was changed every third day. On day 12, cells in NC media were detached with TrypLE and replated on Poly-L-ornithine hydrobromide (Sigma Aldrich) coated 48-well plates at a density of 3,000 cells/well and cultured in NC media.

On day 12, the four lines from each experimental group showing consistent growth were selected from a wider pool of candidate lines. Cell lines were selected following analysis of prior work (Mertens, et al., 2021) with iN lines that showed no substantial sex based differential gene expression (Supplementary Fig.4). On day 20, half the NC media was replaced with CD4^+^ T cell supernatant collected from cultured cells from clinical study participants. On day 21, cells were collected with TrypLE without sorting for either bulk RNAseq or single cell RNAseq.

### Bulk sequencing of iNs

RNA isolation was performed using RNEasy total RNA extraction kit (Qiagen) according to the manufacturer’s instructions. Before library preparation, RNA integrity was assessed by RIN Agilent TapeStation High Sensitivity RNA Screen Tape (Agilent). 50 ng of RNA per sample was poly(A)-selected and amplified (Watchmaker mRNA Library Prep Kit, Watchmaker Genomics, Cat: 7BK0001) and underwent 75 bp paired-end cDNA sequencing (NovaSeq X Plus, Illumina) at the Razavi Newman Integrative Genomics and Bioinformatics Core Facility of the Salk Institute. Transcripts were trimmed to remove sequencing adapters and poly(A) tails (Fastp) (Chen, et al., 2018), then underwent pseudoalignment (Salmon) (Patro, et al., 2017) to the human transcriptome (GENCODE 48).

### Analysis of bulk sequencing data

Protein-coding genes having robust expression across experimental groups were statistically modelled with a negative binomial GLM pipeline (edgeR) (Chen, et al., 2025) in which significances were assessed via likelihood ratio tests. All samples were used for dispersion estimation, and the GLM was formulated for a pairwise contrast between refed and fasted samples from the same participants for each experimental group. The false discovery rate was controlled via the Storey procedure (Storey & Tibshirani, 2003) with π_0_ estimated from the tail of the *p* value distribution. Differences were considered significant when *q* < 0.1.

### Downstream analysis of DEGs

Gene Ontology terms were tested for overrepresentation within upregulated, downregulated, or all regulated genes (Limma: *goana*) (Ritchie, et al., 2015). Transcription factor activities were imputed from the gene expression profiles of each sample (Priori) (Yashar, et al., 2024). Similarly, models of metabolic reaction activity were imputed (Compass) (Wagner, et al., 2021). Gene-set-based derived quantities were considered significant when *p* < 0.01 (gene set overrepresentation: one-sided hypergeometric test; differences in transcription factor activities or in reaction consistencies: paired *t* test). A curated subset of significant GO terms was visualised as a network diagram in clusters based on gene membership (aPEAR; GO terms included: *p* < 0.005, size < 500, and terms relating to other tissue types were manually removed) (Kerseviciute & Gordevicius, 2023).

### Cell-cell communication modelling

Communication was modelled from the gene expression profiles of the CD4^+^ T cells and the secretome-treated iNs (ICELLNET) (Massenet-Regad & Soumelis, 2024). Pathways were highlighted as strongly regulated when there was a large fold change in the communication score between the fasted and re-fed branches of the experiment, *FC* > 3 in either direction.

### Single-cell sequencing of INs

Cell lines treated with disease-matched fasted supernatant and refed supernatant as described above were multiplexed and distributed across two pooled reactions. A total of 20,000 cells per reaction (approximately 2,500 cells per sample) were partitioned using the Chromium GEM-X Single Cell 3′ Chip Kit v4 (10x Genomics, Cat: PN-1000690) and encapsulated with reagents and barcodes from the Chromium GEM-X Single Cell 3′ Kit v4 (10x Genomics, Cat: PN-1000686). Each reaction underwent 90 bp single-end sequencing on a NovaSeq X Plus instrument (Illumina) at the Salk Institute, and reads were aligned and quantified against the GRCh38 reference (10x Genomics Reference 2024-A; GENCODE 44) with Cell Ranger 9.0.1 using *--include-introns*. Pooled cells were genetically demultiplexed to their donor of origin using polymorphism data from prior gDNA sequencing of each cell line (Demuxlet) (Kang, et al., 2018). Ambient RNA was modelled, cell barcodes associated with only ambient RNA were removed from the analysis, and transcript counts adjusted for background subtraction were imputed for each cell (CellBender) (Fleming, et al., 2023). Droplets predicted to contain doublets (via Demuxlet) or only ambient RNA (via CellBender) were filtered from downstream analysis.

### Analysis of single-cell sequencing data

Transcript counts for each cell were imported into R (Seurat) and the data were harmonised across reactions (Seurat: *SCTransform*) (Hao, et al., 2024). Clusters based on nearest neighbour gene expression were labelled according to classical marker genes and differential marker genes between subsets of cells were discovered via fold change and *p* value thresholds, |log2 FC| > 0.1 and *p* < 0.01 (Wilcoxon rank sum test). Gene Ontology terms were tested for upwards, downwards, or mixed-direction enrichment within the set of differential marker genes scored via their directional log_10_ *p* value (Limma: *cameraPR*) (Ritchie, et al., 2015) and were considered significant when *p* < 0.01 (modified Wilcoxon rank sum test accounting for a fixed inter-gene correlation *c* = 0.01).

### Metabolomics of CD4+ T cell supernatant

A sample of each pooled supernatant treatment was using modifications of previously described methods (Bligh & Dyer, 1959). A mixture of deuterium-labeled internal standards (Sigma-Aldrich) were added to the media samples at the start of the extraction procedure, followed by water, methanol, and chloroform. Upon vortexing and centrifugation, the aqueous layer was separated and dried, reconstituted in 20% methanol (100 μL), vortexed, and transferred to HPLC vials for LC-MS analysis.

Untargeted metabolomic analysis of polar small molecules was performed on a Thermo Scientific Vanquish UHPLC coupled to a Q-Exactive quadrupole-orbitrap mass spectrometer (Thermo Fisher Scientific) using modifications of a previously published LC method (Larios-Serrato, et al., 2025). Mass spectra were acquired under both positive and negative ionization, using injection volumes of 2 and 5 μL, respectively (sheath gas flow rate: 55; aux gas flow rate: 10; sweep gas flow rate: 2; ion transfer tube temperature: 275°C; vaporizer temperature: 320°C). Spray voltage was 3.5 and −2.6 kV for positive and negative ionization, respectively. Full scan mass spectrum resolution was set to 60,000 with a scan range of *m*/*z* 60 to 900. For MS/MS scan, stepped collision energies (%) of 20, 35, 50 were used and the mass spectrum resolution was set to 15,000.

All processing of the LC-MS raw data files were performed using MS-DIAL 5.5 software for data collection, peak detection, alignment, adduct, and identification (Tsugawa, et al., 2015). Compounds were annotated by *m*/*z*, MS/MS spectra using the libraries’ MassBank of North America (MoNA) and Global Natural Products Social Molecular Networking (GNPS). Internal standards were monitored for retention time and intensity and PCA was used for multivariate statistics and visualization, specifically for outlier detection. From the MS-DIAL results file, all detected features/metabolites were removed if *sample max* / *blank average* < 10. Known (positively identified/annotated) features/metabolites were manually evaluated to confirm accuracy. With regard to detected features not annotated/identified (unknown compounds), following removal based on the previously mentioned *sample max* / *blank average*, only those with a *m*/*z* and MS2 data were retained and are reported. For quality control purposes, pooled QC project samples were run to assess reproducibility, external pooled plasma samples (BioIVT) were run to compare to historical method performance, a method blank was used to assess background, solvent blanks were run throughout the sequence to confirm the absence of carryover, and samples were analyzed in randomized order.

### Lipidomics of CD4^+^ T cell supernatant

A mixture of 69 deuterium-labeled internal standards (UltimateSPLASH ONE, Avanti Polar Lipids) was added to the supernatant samples at the start of the extraction procedure, followed by water, methanol, and chloroform. Upon vortexing and centrifugation, the organic layer was separated and dried, reconstituted in 9:1 methanol:toluene (100 μL), vortexed, and transferred to HPLC vials for LC-MS analysis.

Lipidomic analysis was performed on a Thermo Scientific Vanquish UHPLC coupled to a Q-Exactive quadrupole-orbitrap mass spectrometer (Thermo Fisher Scientific) using previously disclosed protocols (Beaufrère, et al, 2025). Lipids were separated using a Waters Acquity BEH C18 column (50 mm × 2.1 mm; 1.7 μm) with an Acquity BEH C18 VanGuard pre-column (5 mm × 2.1 mm; 1.7 μm) (Waters). The column temperature was 65 °C. LC-MS analysis was performed at a flow rate of 0.8 mL/min with a gradient ranging from start conditions of 15% B to 99% B. Mobile phase A consisted 60:40 acetonitrile:water and mobile phase B consisted 90:10 isopropanol:acetonitrile. For optimal detection of lipids, different mobile phase modifiers were used for positive and negative ionization mode. Positive ionization mode contained 10 mM ammonium formate and 0.1% formic acid, while negative ionization mode contained 10 mM ammonium acetate. An injection volume of 2 and 5 μL were used for positive and negative ionization modes, respectively.

All processing of the LC-MS raw data files were performed using MS-DIAL 5.5 software for data collection, peak detection, alignment, adduct, and identification (Tsugawa, et al., 2015). The detailed parameter setting was as follows: MS1 tolerance, 0.01 Da; MS2 tolerance, 0.025 Da; minimum peak height, 100,000 amplitude; mass slice width, 0.05 Da; smoothing method, linear weighted moving average; smoothing level, 3 scans; minimum peak width, 5 scans. [M + H]^+^, [M + NH_4_]^+^, [M + Na]^+^, [2M + H]^+^, [M + H - H_2_O]^+^ and [M - H]^−^, [M - 2H]^2-^, [M + CH_3_OO]^−^, [M - H - H_2_O]^−^ were included in adduct ion setting for positive and negative mode, respectively. Compounds were annotated by *m*/*z* and MS/MS spectra against the LipidBlast mass spectra database (Kind, et al., 2013). Internal standards were monitored for retention time and intensity and PCA was used for multivariate statistics and visualization, specifically for outlier detection. From the MS-DIAL results file, all detected features/metabolites were removed if *sample max* / *blank average* < 10. Known (positively identified/annotated) features/metabolites were manually evaluated to confirm accuracy. With regard to detected features not annotated/identified (unknown compounds), following removal based on the previously mentioned sample max/blank average, only those with a *m*/*z* and MS2 data were retained and are reported. For quality control purposes, pooled QC project samples were run to assess reproducibility, external pooled plasma samples (BioIVT) were run to compare to historical method performance, a method blank was used to assess background, solvent blanks were run throughout the sequence to confirm the absence of carryover, and samples were analyzed in randomized order.

### Proteomics of CD4^+^ T cell supernatant

Proteins in conditioned media were precipitated using a chloroform-methanol-water method. The proteins were then resuspended in 8 M urea in 50 mM triethylammonium bicarbonate (TEAB). Cysteine disulfide bonds were reduced with 10 mM TCEP for 30 minutes and alkylated with 40 mM CAA for 1 hour in the dark at room temperature. The urea concentration was then diluted to < 1 M with additional 50 mM TEAB, and the proteins were digested overnight with 500 ng of trypsin at 37 C (Pierce). The digestion was stopped with 1% TFA, and the peptides were desalted with C18 Spin Tips (Pierce). Peptide amounts were normalized using a colorimetric peptide assay (Pierce).

After drying in a SpeedVac, the remaining peptides were resuspended in 0.1% FA in 5% ACN and approximately 400 ng were injected into a Thermo Neo Vanquish coupled to an Orbitrap Eclipse Tribrid mass spectrometer. Peptides were separated on a 50 cm Acclaim PepMap C18 column (Thermo Fisher) at a flow rate of 300 nL/min. The mobile phases were 0.1% FA in water (A) and 0.1% FA in 80% ACN (B), and the gradient started at 4% B before going up to 30% B over 120 minutes, followed by an increase to 50% B over 10 minutes. The column was then washed at 90% B and re-equilibrated for the next sample. The mass spectrometer was operated in positive mode with data-dependent acquisition. Full MS scans from 375–1500 *m*/*z* were acquired in the Orbitrap at 120k resolution. The most abundant precursors from each MS1 scan were subjected to HCD fragmentation (30 NCE), and the resulting MS2 scans were collected in the Orbitrap at 30k resolution over a total cycle time of 3 seconds. The isolation window was 0.7 m/z, and the dynamic exclusion time was 30 seconds.

LC-MS data were analyzed with Proteome Discoverer v3.2 using the Sequest search engine against the UniProt *Homo sapiens* reference database (83,078 entries; downloaded March 2025). Sequest settings included a precursor mass tolerance of 10 ppm and a fragment mass tolerance of 0.02 Da. Tryptic digestion was specified with a maximum of 2 missed cleavages. Modifications included fixed cysteine carbamidomethylation and variable methionine oxidation and N-terminal acetylation. Protein identifications were filtered with Percolator according to a false discovery rate (FDR) cut-off of 1%. Label-free protein quantitation was done using the Minora feature detector and precursor quantifier nodes in ProteomeDiscoverer. Protein abundances were normalized by total peptide amount. Peptides were required to be in at least 50% of the replicates to be summed and used for calculation of protein abundance.

## Acknowledgements

We are grateful to all donors who participated. We acknowledge the Joint Clinical Research Facility, ILS, Swansea University for their support with the clinical study. We acknowledge the support of the Supercomputing Wales project, which is part-funded by the European Regional Development Fund (ERDF) via Welsh Government. This work was supported by the Mass Spectrometry Core of the Salk Institute (RRID:SCR_014843) with funding from NIH-NCI CCSG P30 CA014195, NIH-NIA San Diego Nathan Shock Center P30 AG068635, an NIH S10 award for metabolic instrumentation S10 OD021815, and the Helmsley Center for Genomic Medicine. This work was supported by the Razavi Newman Integrative Genomics and Bioinformatics Core Facility of the Salk Institute (RRID:SCR_014842 and SCR_014846) with funding from NIH-NCI CCSG P30 CA014195, NIH-NIA San Diego Nathan Shock Center P30 AG068635, the NIH-NIA Liver Cancer P01 AG073084-04, the Howard and Maryam Newman Family Foundation and the Helmsley Trust. PCD is supported by the Medical Research Council (grant no. MR/W000229/1). PCD and AJY are supported by the NIHR Newcastle Biomedical Research Centre awarded to the Newcastle upon Tyne Hospitals NHS Foundation Trust, Newcastle University and Cumbria, Northumberland, Tyne and Wear Foundation Trust. This work was supported by grants from the Hilary & Galen Weston Foundation (grant no. UB190138), BRACE, the Alzheimer’s Charity (grant no. BR2286), Coleg Cymraeg Cenedlaethol (grant no. YSG22/02) and a US-UK Fulbright Scholar Award to JSD. We thank Mary Lynn Gage and Jessica Stoneman for editorial comments.

---

**Supplementary Fig. 1:**
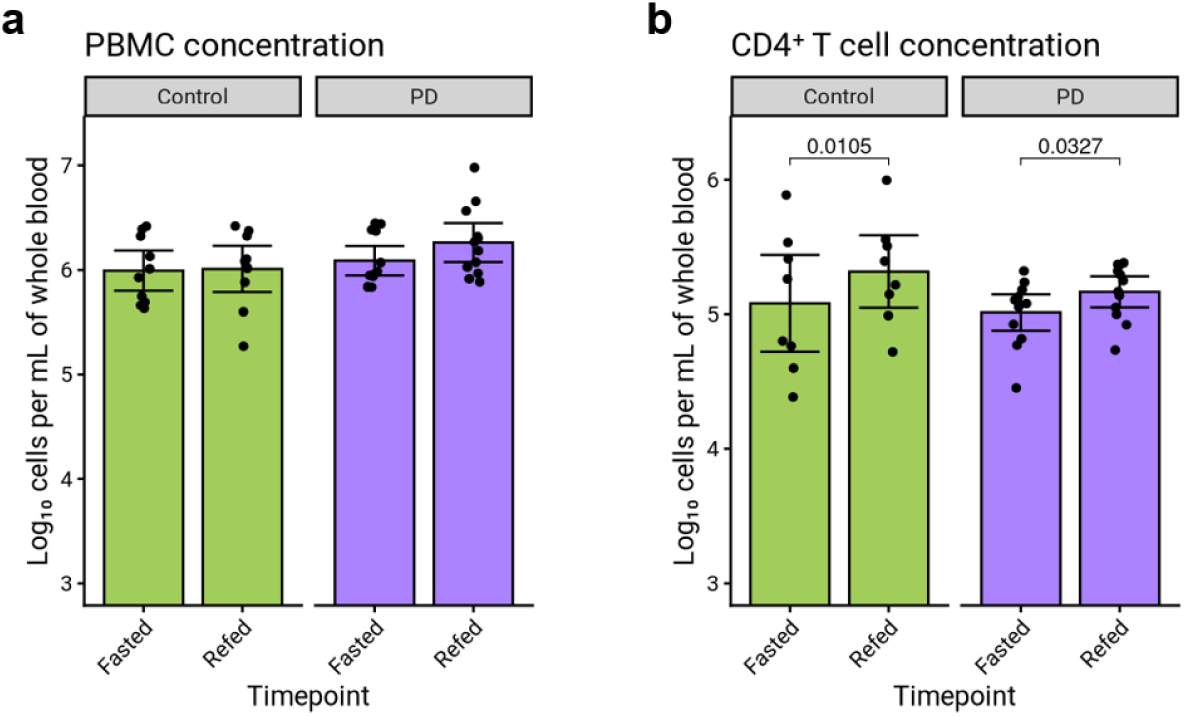
Abundance of PBMCs and CD4^+^ T cells in donated blood. **a**, Comparison of the measured concentration of PBMCs in whole blood between the refed and fasted timepoints for healthy control participants (*n* = 10) and PD participants (*n* = 12) showing no statistically significant change; paired *t* tests, *p* < 0.05. **b**, The measured concentration of CD4^+^ T cells in whole blood increased significantly in both PBMCs (*n* = 8; *p* = 0.0105) and CD4+ T cells (*n* = 8; *p* = 0.0327).

**Supplementary Fig. 2:**
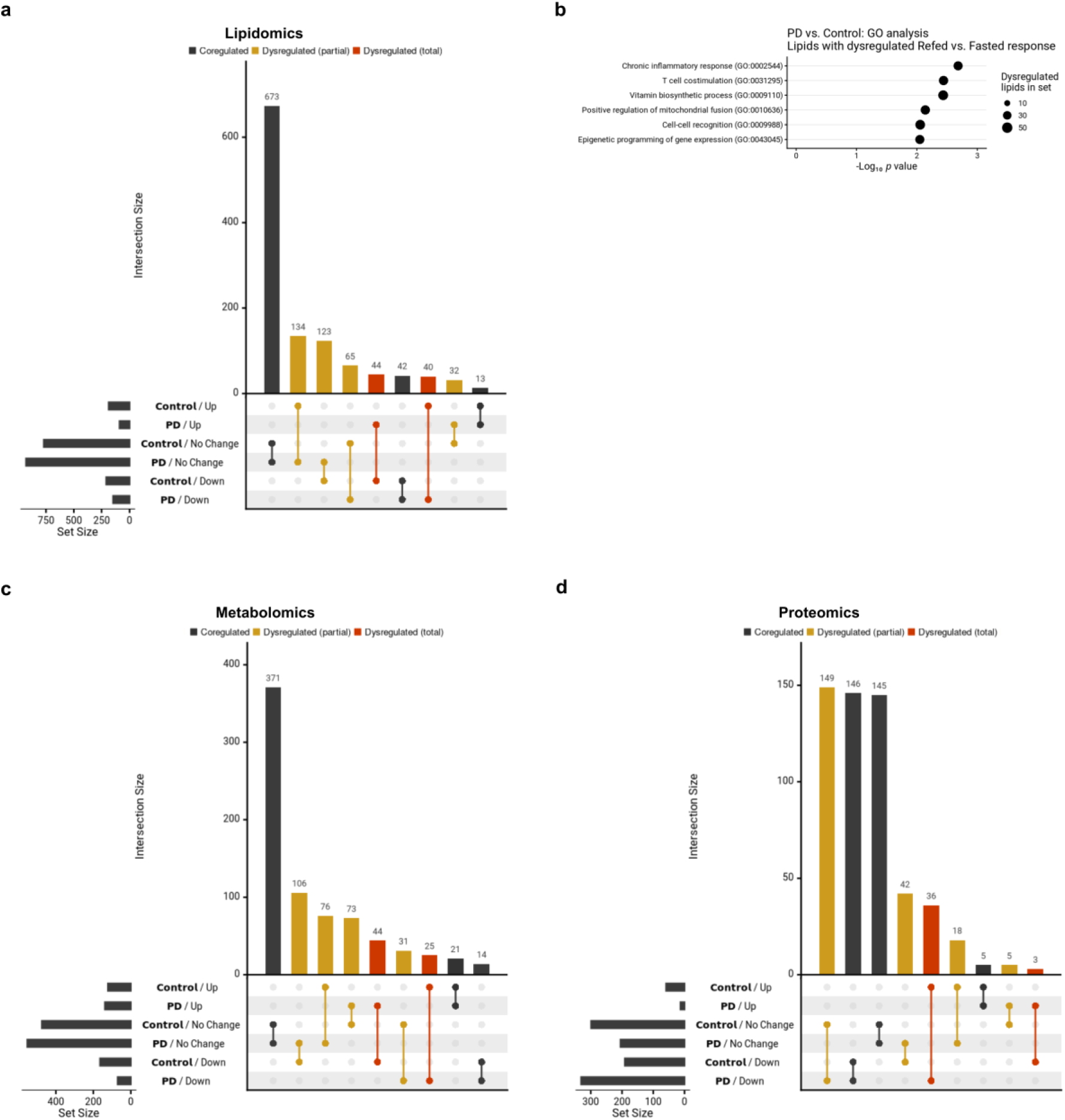
Chemical content of pooled supernatant treatments. **a**, UpSet plot categorising lipids detected in the pooled CD4^+^ T cell supernatant treatment according to their fasting-refeeding regulation in both the healthy control and PD disease conditions; analytes were called as upregulated or downregulated in a disease condition if they had a fold change > 1.5 in that direction between the refed and fasted sample, black: subsets which were regulated in the same direction in both disease conditions, yellow: subsets not regulated in one condition and regulated in the other, red: subsets regulated in opposite directions in the two conditions. **b**, Selected terms from a Gene Ontology overrepresentation analysis of dysregulated lipids: pathways implicate inflammation and metabolism; MAPtoGO analysis with background set to detected lipids, *p* < 0.01. **c**, UpSet plot categorising detected small molecule metabolites as per **a**. **d**, UpSet plot categorising detected proteins as per **a**.

**Supplementary Fig. 3:**
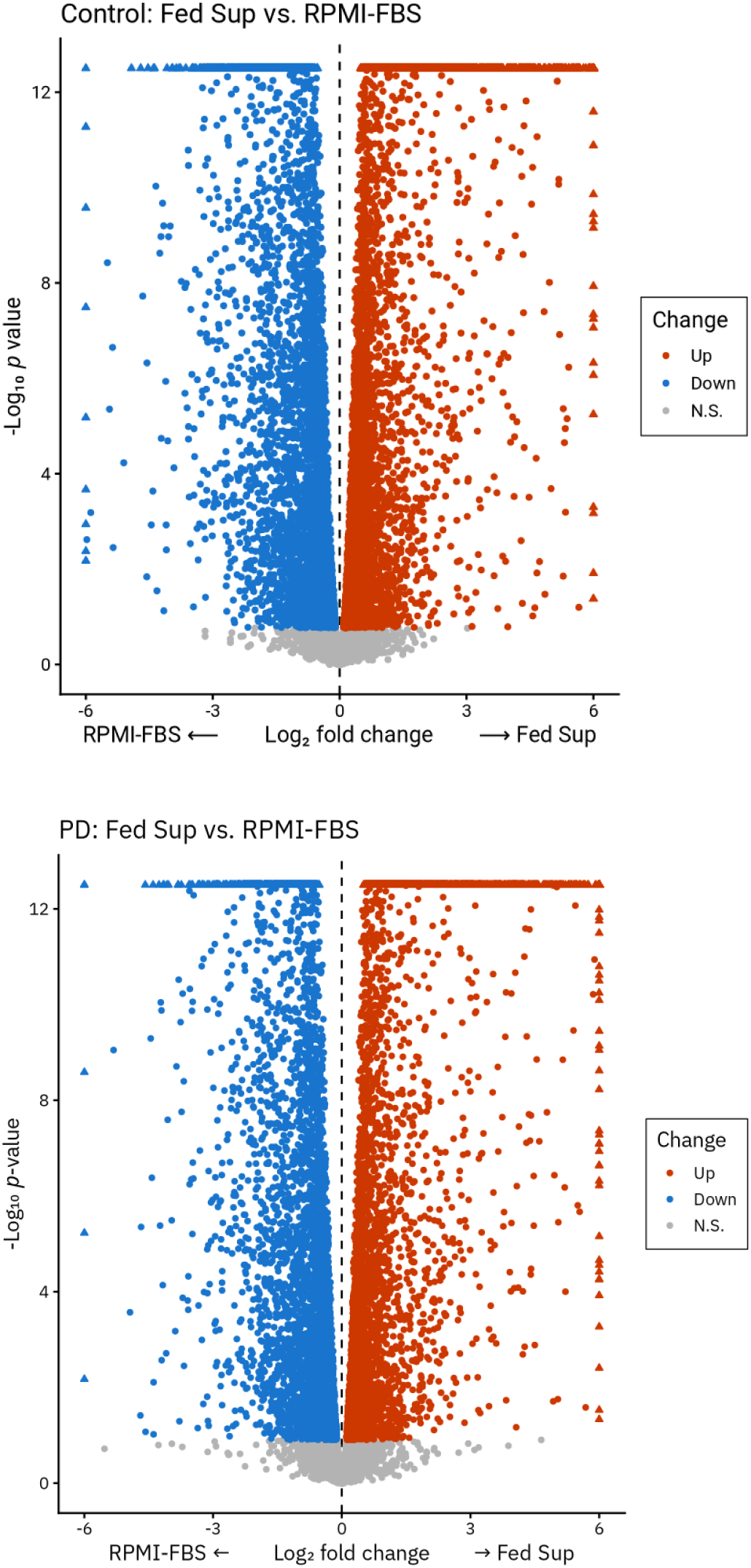
Response of iNs to a pooled supernatant treatment. Differential gene expression analysis showing 10309 DEGs when comparing refed CD4^+^ T cell supernatant treatment to RPMI-FBS vehicle media treatment in healthy control iNs (*n* = 4) and 7754 DEGs in PD iNs (*n* = 4), triangles capped for plotting; edgeR, likelihood ratio tests adjusted with Storey’s procedure, *q* < 0.1.

**Supplementary Fig. 4:**
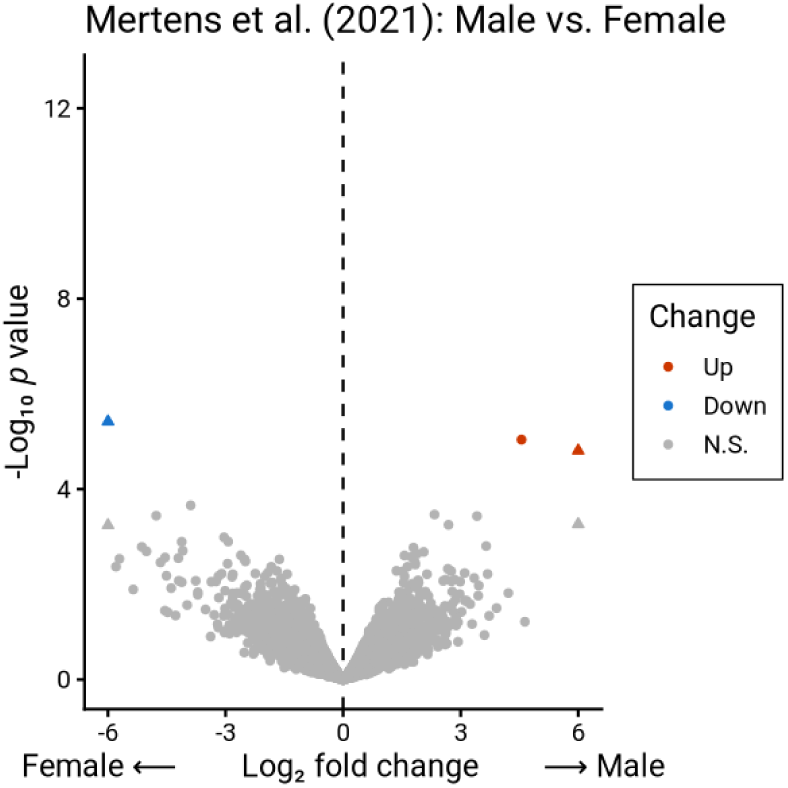
No evidence for substantial sex differences in gene expression of control iN lines from Mertens, et al. (2021) Differential gene expression analysis showing 3 DEGs when comparing healthy control iN lines from male (*n* = 3) and female (*n* = 3) donors, triangles capped for plotting; edgeR, likelihood ratio tests adjusted with Storey’s procedure, *q* < 0.1. Raw sequencing data from Mertens, et al. (2021), ArrayExpress: E-MTAB-10352.

**Supplementary Table 1:**
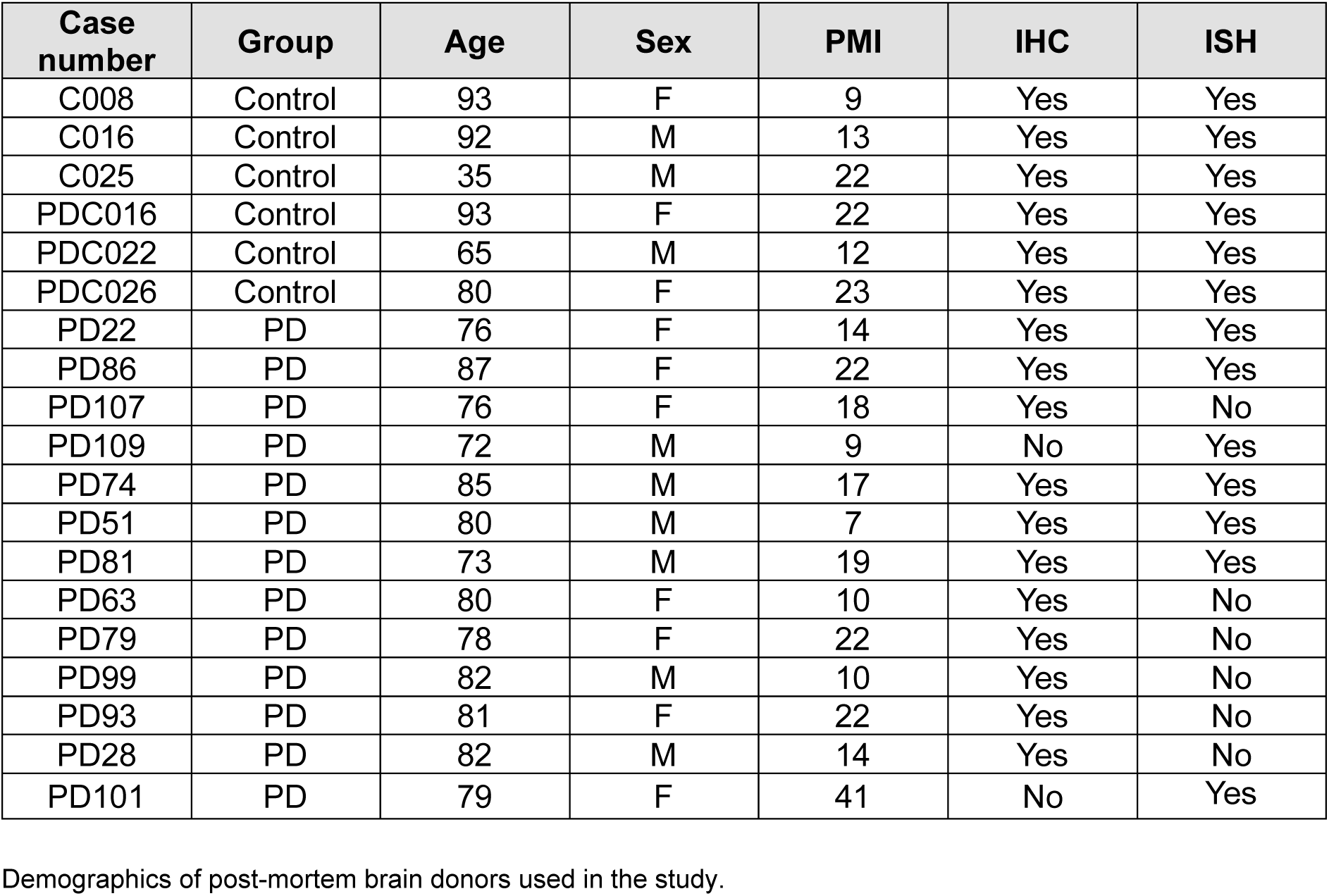
Post-mortem brain tissue donor demographics.

**Supplementary Table 2:** Blood donor demographics.

| Donor number | Group | Sex | Age | Immune cell abundance analysis | Cytokine panel analysis | Pooled secretome treatments | RNAseq analysis |
| --- | --- | --- | --- | --- | --- | --- | --- |
| S1 | Control | Male | 73 | Yes | Yes | Yes | No |
| S2 | Control | Male | 67 | Yes | Yes | Yes | Yes |
| S3 | PD | Male | 80 | Yes | No | No | No |
| S6 | PD | Male | 67 | Yes | Yes | Yes | No |
| S7 | PD | Male | 75 | Yes | Yes | Yes | Yes |
| S8 | PD | Female | 77 | Yes | No | Yes | No |
| S9 | PD | Female | 73 | Yes | Yes | Yes | Yes |
| S10 | PD | Female | 70 | Yes | No | Yes | No |
| S12 | Control | Female | 65 | Yes | Yes | Yes | Yes |
| S14 | PD | Male | 61 | Yes | No | Yes | No |
| S16 | PD | Female | 74 | Yes | No | Yes | Yes |
| S17 | PD | Male | 70 | Yes | Yes | Yes | Yes |
| S18 | Control | Female | 71 | Yes | Yes | Yes | Yes |
| S19 | PD | Female | 72 | Yes | No | No | No |
| S20 | Control | Male | 73 | Yes | No | No | No |
| S23 | Control | Female | 78 | Yes | Yes | Yes | Yes |
| S24 | Control | Male | 79 | Yes | Yes | Yes | Yes |
| S25 | Control | Male | 77 | Yes | Yes | Yes | No |
| S27 | Control | Female | 61 | Yes | Yes | Yes | No |
| S29 | Control | Female | 62 | Yes | Yes | Yes | No |
| S30 | PD | Female | 60 | Yes | Yes | Yes | Yes |
| S34 | PD | Male | 81 | Yes | No | Yes | No |

Demographics of participants that donated blood used in the study.

**Supplementary Table 3:** Pooled supernatant treatments.

| Group | Timepoint | Donor numbers |
| --- | --- | --- |
| Control | Fasted | S1, S2, S12, S18, S23, S24, S25, S27, S29 |
| Control | Refed | S2, S12, S18, S23, S24, S25, S27, S29 |
| PD | Fasted | S6, S7, S8, S9, S10, S14, S16, S17, S30, S34 |
| PD | Refed | S6, S7, S8, S9, S10, S14, S16, S17, S30, S34 |

Details of pooled samples from study participants used in the pooled secretome treatments.

**Supplementary Table 4:**
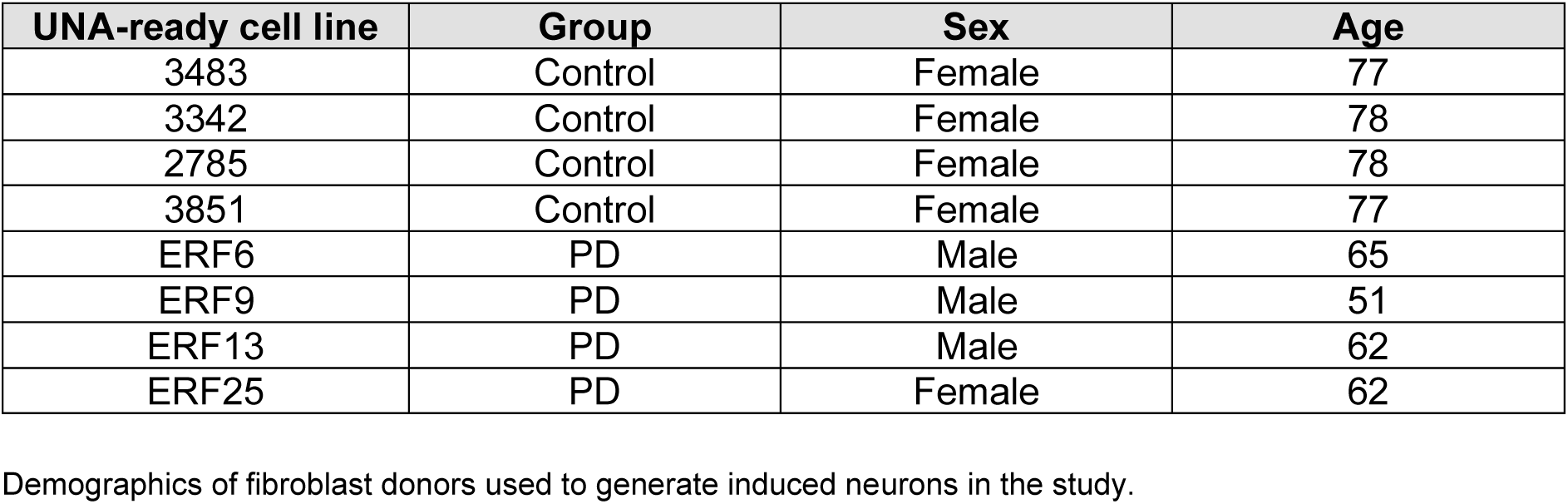
iN line demographics.

